# Discrete-time quantum walks reveal critical residues involved in long-range interactions in proteins

**DOI:** 10.64898/2026.10.26.761941

**Authors:** Ramnarayan Bera, Swati Sharma, Deep Nath, Madhurima Khamaru, Devrani Mitra, Soumen Roy

## Abstract

Long-range interactions between residues in proteins underlie diverse biological processes such as signal transmission, conformational regulation, and catalysis. Identifying residues that mediate such communication remains a major challenge. Herein, we introduce a framework based on quantum walks and entropy, and employ it on protein structures to predict important residues participating in long-range interactions. The selected proteins have been well-studied experimentally in literature. Our approach models information flow through protein contact networks using discrete-time quantum walk dynamics, where quantum superposition and interference govern transition amplitudes at the residue level. Normalized time-averaged transition probabilities quantify interaction strength between residues, while Shannon entropy calculations identify residues exhibiting delocalized interaction behavior. We find that the residues identified by us are known to be structurally and functionally important in literature, thereby confirming the predictive capacity of this framework. We experimentally validate our formalism on *As*LOV2 (PDB: 2V0U) using site-directed mutagenesis, protein purification, and dark-recovery kinetics. The predicted residues are important for understanding the mechanism of photosensing in *As*LOV2.

**Significance:** This study bridges quantum transport theory and structural biology by introducing discrete-time quantum walks as a mechanism to model communication between residues. We provide a scalable and interpretable method to predict key residues, without requiring molecular dynamics or prior functional annotation. The present method, inspired by quantum physics and quantum information, and validated by experiments, offers a broad and robust framework to study communication in biomolecular systems. It further demonstrates how quantum principles such as coherence and interference can enhance our understanding of information flow in biological macromolecules.

## I. INTRODUCTION

Proteins are chains of amino acids, folded into specific three-dimensional structures stabilized by a combination of covalent and non-covalent interactions [1]. It is well established that the structure and function of a protein are closely correlated. Even subtle local structural perturbations alter residue–residue interaction patterns, which correspond to different conformational states and biochemical activities [2]. While short-range interactions are often conserved across protein conformations, variations in long-range interactions lead to the emergence of distinct functional states [3, 4]. Long-range interactions play a distinct role in determining the tertiary structure of a protein, while short-range interactions largely contribute to secondary structure formation [5]. These long-range couplings define the main communication pathways across the protein, guiding conformational changes and functional responses upon ligand binding. Thus, in the present work, both long-range interactions and long-range communication pathways are considered equivalent descriptors of information flow within proteins.

The interactions among amino acid residues can be represented as a protein contact network (PCN), where residues act as nodes and the edges represent the interactions between them [6, 7]. Since proteins possess many degrees of freedom(when we consider the whole atomic details), analyzing PCNs offers a coarse-grained yet insightful approach to understand protein function without explicitly modeling atomic details unlike computationally expensive methods (e.g., MD-based simulation).

Understanding and identifying residues that mediate long-range communication is fundamentally important for multiple reasons. First, such residues often act as allosteric communication hubs, transmitting perturbations from ligand-binding sites to distant functional domains and thus serving as critical determinants of allosteric regulation [4, 8]. Second, these residues are frequent targets for natural mutations associated with disease, making their identification crucial for understanding disease mechanisms and for rational design of therapeutics [9]. Third, identifying such residues illuminates fundamental principles of how proteins evolve regulation and switch between functional states, informing both evolutionary biology and synthetic biology approaches.

A variety of computational strategies (i.e., network-based, Molecular dynamics (MD)-based, coevolution and sequence-based approaches etc.) have been developed to pinpoint residues important for long-range communication. Yet, each approach carries its own limitations i.e., computationally expensive (MD-based), data-demanding (sequence-based), or theoretically limited to diffusive, Markovian assumptions (classical network methods).

We have developed a discrete-time quantum walk (DTQW) and entropy-based framework that captures coherent information propagation without requiring evolutionary information for identifying long-range interaction residues. In our framework, a quantum walker coherently traverses a PCN with unitary evolution governed by a coin-shift operator scheme. By computing the DTQW time-averaged transition probability (TATP) from each residue to all others, we obtain an effective communication strength between residue pairs that reflects both network connectivity and quantum-mechanical interference effects. Shannon entropy of the normalized TATP distribution for each residue quantifies the distributional diversity or “broadness” of a residue’s communication reach. A high-entropy residue distributes its outgoing communication broadly across many targets, indicating it acts as a hub of communication to spread probability amplitude that reaches diverse regions of the PCN. Residues exhibiting both elevated transition probability (strong communication strength) and high Shannon entropy (broad, distributed communication) are designated as critical residues for long-range communication.

Our approach operates directly on static structural data from Protein Data Bank (PDB) without requiring dynamical simulations of structure, large sequence datasets or biochemical priors. It can be deployed on any protein structure, including those from cryo-EM or AlphaFold predictions, making it suitable for high-throughput screening, drug discovery, and protein engineering efforts. The integration of Shannon entropy provides an information-theoretic lens that is interpretable, computable, and invariant to network size and scaling. Quantum walks on networks have been extensively studied in physics and mathematics, providing a rigorous, well-characterized framework with clear mathematical properties and limiting behaviors [10, 11].

The combination of coherence-based transport (leads to quantum superposition and interference) and entropy-based analysis provides a novel, quantitative perspective. This framework may offer new insights into the fundamental principles underlying allosteric regulation and long-range signaling in biomolecules, with practical applications in predicting allosteric sites, guiding protein engineering, and informing structure-based drug discovery targeting allosteric modulators.

To verify the predictive power of the disclosed framework, we not only surveyed the literature for the experimental system, but also verified it experimentally. We validate our findings on a well-studied light-sensing LOV2 domain of *Avena sativa* phototropin 1 (AsLOV2, PDB ID: 2V0U) [12]. This system served as a primary benchmark for correlating the theoretical quantum walk dynamics with observed biological phenomena, following the residue-level theory-and-experiment strategy established for LOV photoreceptors [13, 14]. Phototropins are blue-light-activated serine/threonine kinases found in plants and algae that not only regulate the highly important phenomenon seen in plants — phototropism (bending of plants towards light) — but also chloroplast relocation and stomatal opening [15]. Each phototropin contains two light-oxygen-voltage (LOV) sensing domains (LOV1 and LOV2), and a C-terminal serine-threonine kinase domain [16]. Upon absorption of blue light (approximately 450 nm), each LOV domain forms a reversible covalent adduct between a conserved cysteine residue and the C(4a) atom of its non-covalently bound flavin mononucleotide (FMN) cofactor [16, 17].

The LOV2 domain of *Avena sativa* phototropin 1 (AsLOV2) has become the most widely exploited optogenetic actuator in the field [18, 19]. In the dark state, a C-terminal amphipathic helix designated the J*α*-helix is docked against the LOV *β*-scaffold surface through a network of hydrophobic contacts and hydrogen bonds [12, 20]. Upon blue-light illumination, the formation of the cysteinyl-FMN adduct triggers a conformational change that propagates to the J*α*-helix, causing it to undock from the *β*-scaffold and unfold [20, 21]. Signal propagation from the chromophore pocket to the peripheral helix proceeds through a conserved glutamine on the I*β* strand, with a secondary route involving the N-terminal A^′^*α* helix [22, 23].

A fundamental understanding at the molecular level is the primary requirement to utilizing the protein in biotechnological applications [24]. For example, AsPHOT1 LOV2 has been widely used in developing LOV-based optogenetic designs [18, 19, 25]. More insights at the residue level can help improve such designs [24, 26]. Moreover, phototropin LOV2-based fluorescent reporter proteins have recently revolutionised imaging in cell biology [27–29] These reporters hold a selective advantage over the established fluorescence reporter GFP in terms of (A) lower molecular weight, approximately 10 kDa versus approximately 25 kDa for GFP [28], and (B) their ability to thrive in both aerobic and anaerobic conditions, since flavin-based fluorescence requires no oxygen-dependent chromophore maturation [27]. Therefore, unravelling the residue-level information can significantly facilitate LOV-based designs in both optogenetics and imaging technologies. Hence, these structural transitions at the molecular level have been harnessed to engineer photoactivatable proteins [18], light-gated protein-protein interaction switches [19, 25], and optogenetic tools in which a protein of interest is fused to or inserted into the AsLOV2 domain, thereby gating its activity by illumination [30, 31].

The residues identified by the computational analyses of AsLOV2 in its dark state (PDB: 2V0U) are THR417, ASN418, LEU496, ALA497, ASN508, PHE509, LEU510, and GLN511, which are located within the J*α*-helix and at its interface with the LOV *β*-scaffold, as participating in contacts that stabilize the docked conformation of the J*α*-helix [12, 20, 22].

## II. RESULTS AND DISCUSSION

We applied our framework based upon discrete-time quantum walk (DTQW) and entropy, to a set of well-studied proteins listed in Table I. Because these proteins have been extensively characterised experimentally, with known allosteric sites, catalytic residues, and functional hotspots, we confirmed the functional importance of the predicted residues by comparing them with the existing literature. In addition, we experimentally validated our predictions for *As*LOV2 by measuring FMN binding through dark- and light-state kinetics assays of the wild-type protein and its mutants. Here, we present the complete analysis for eight representative proteins.

**TABLE I:** Important Residues Identified from DTQW-Entropy Analysis.

| PDB ID | Important Residues |
| --- | --- |
| 5A8B | ASN251, PHE252, VAL253, VAL254, PHE289, LEU290, GLN291, MET313, SER314, VAL315, ASN329, GLN330, PHE332, ILE333, GLY348, VAL349, GLN350, CYS351, LYS352 |
| 4DJA | PRO261, ASN262, LEU263, HIS265, LEU267, LEU268, SER269, PHE270, ILE272, ASN273, GLY275, LEU277, ARG311, GLU312, MET314, ARG315, ASP324, TYR325, VAL326, GLU357, THR358, TYR363, ALA362, HIS365, HIS366, GLN368, LEU370, GLY374, LYS385, HIS388, VAL394, ALA396, ASP397, TYR399, VAL402, LEU404, ILE408, SER411, ASP415 |
| 3ZXS | PRO257, PHE258, SER265, SER266, MET268, ASN269, PRO275, ARG307, GLU308, TYR321, ILE322, ARG323, SER324, GLN353, LEU357, ALA358, TYR359, HIS361, HIS362, LEU366, GLY370, ASN371, PRO380, HIS384, VAL390, ILE392, ASP393, LEU395, TRP397, VAL398, ALA400, ILE404, GLY405, MET406, SER407 |
| 8Z1J | LEU208, THR209, VAL210, PHE211, ALA218, ARG221, LEU222, PRO248, ALA249, THR250, THR251, VAL252, LEU253, SER254, PRO255, LEU262, LEU349, VAL350, THR351, TRP352, GLY353, TRP354, HIS356, HIS361, HIS386, ILE388, ASP389, LEU402 |
| 9HNM | ARG258, PRO260, SER268, PRO269, LEU271, ASN272, PRO278, ARG310, GLU311, ALA325, GLN326, ARG327, ASP352, ARG355, ALA356, HIS360, ALA361, HIS365, LEU369, GLY373, PRO383, ASP384, VAL393, ASP396, ARG407, ALA410, THR411 |
| 1MDN | PHE8, LEU9, VAL10, VAL11, ASP12, PHE53, VAL54, ILE55, SER56, ASP57, TRP58, LEU81, PRO82, VAL83, MET85, VAL86, SER104, GLY105 |
| 1F4V | PHE8, LEU9, VAL10, VAL11, ASP12, PHE53, VAL54, ILE55, SER56, ASP57, TRP58, LEU81, PRO82, VAL83, MET85, VAL86, SER104, GLY105 |
| 2V0U | LYS413, ASN414, PHE415, ILE417, THR418, PHE429, ALA430, SER431, PHE494, HIS495, LEU496, GLN497, TYR508, ILE510, GLY511, VAL512, GLN513 |

### A. Proposed Method

#### 1. DTQW Dynamics on Protein Contact Networks

To investigate long-range communication residues within proteins, our formalism was applied to protein contact networks (PCNs) constructed from residue-residue interaction maps derived from crystallographic structures at a 4.5 Å contact cutoff [32, 33].

For each protein, the Hilbert space was defined as

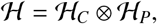

where *H*_*P*_ corresponds to the residue positions and *H*_*C*_ encodes local directional degrees of freedom towards the residue’s all nearest neighbors. The walker’s state was initialized at each residue with an equal mixture over all outgoing coin states, ensuring unbiased probing of the local topological environment.

At each time step, the quantum state evolved under the unitary operator

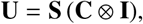

where **C** is the coin operator (Fourier or Grover) and **S** implements the graph-dependent shift. The transition probability from initial residue *i*_0_ to residue *j* at time *t* is computed as

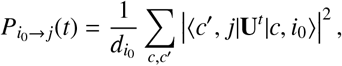

encoding interference patterns generated by coherent spreading of the walker. The normalization by degree 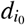 accounts for the number of outgoing edges and ensures proper probability accounting from each starting residue.

Quantum interference leads to temporal oscillations in 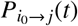, contrasting sharply with the monotonic diffusion characteristic of classical random walks. To obtain a stationary and interpretable measure of communication between residues, we computed the *time-averaged transition probability* (TATP),

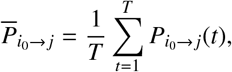

where *T* = 100 time steps was chosen to ensure convergence while remaining computationally efficient. This time averaging removes short-term quantum oscillations while retaining the global transport characteristics induced by network topology. The resulting TATP matrix 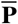 becomes approximately doubly stochastic for sufficiently large *T*, indicating balanced probability flow across the protein and confirming the unitary, non-dissipative nature of the dynamics.

We tested our framework for two different coin operators to assess the sensitivity of our framework introducing directional bias to a initial state of equal mixtures of all outgoing directions:

- **Fourier coin:** Produced broader and more symmetric probability propagation, leading to higher delocalization and more homogeneous spreading across residues. This behavior makes the Fourier coin especially sensitive to identifying regions involved in collective or long-range communication pathways.
- **Grover coin:** Produced more localized spreading with enhanced concentration on nearby residues, highlighting local communication hubs and densely connected regions.

The choice of coin operator leads to almost same critical residues by setting different cut-off/threshold as the involved length scales of Shannon entropies of all the residues are different under different coin operators. For the analysis presented here, we primarily employed the Fourier coin, which proved most effective at identifying experimentally known critical residues.

#### 2. Identifying Important Residues for Long-Range Communication

The DTQW-derived transition matrix generally provides a quantitative picture of how quantum perturbations propagate through the given networks. To identify residues actively participating in long-range interactions and communication, we performed a multi-step analysis of the time-averaged probability distributions (TATP).

a. *Step 1: Normalization and Probability Distributions*. For each residue *i*_0_, the time-averaged transition probabilities were normalized to obtain a probability distribution,

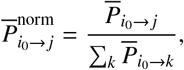

which describes the spatial extent and pattern of perturbation spreading originating from residue *i*_0_. This normalization enables us to compare a broad connectivity of each residue with others. Representative normalized time-averaged transition probability profiles for four well-characterized proteins (1F4V: CheY, 2V0U: *As*LOV2, 1MDN: Myoglobin, and 5A8B: *Pt*Aureo1a LOV) are displayed in FIG. 1. The heatmaps reveal distinct communication patterns: some residues exhibit tightly localized spreading (appearing as bright diagonal spots), while others show delocalized, broad propagation patterns. These communication patterns encode the information of residues involved in long-range interaction. The following steps are needed to decode that information.
b. *Step 2: Shannon Entropy as a Perturbation-Flow Descriptor*. The Shannon entropy of residue *i*,

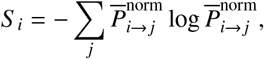

quantifies the diversity and uncertainty of different transitions emanating from residue *i*. Entropy values range from 0 (deterministic spreading to a single residue) to ln(*N*) (uniform spreading across all *N* residues). For typical proteins with *N* ≈ 200-400 residues, maximum entropy is ln(400) ≈ 5.99. To avoid numerical issues with zero probabilities, we set 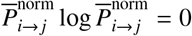 when 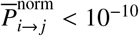. Mathematically, entropy measures the extent to which probability is spread across the available residue targets:

- **Low entropy:** Indicates that residue *i* concentrates its communication probability on a few specific targets,structurally constrained residues with limited outgoing interactions.
- **High entropy :** Indicates that residue *i* distributes quantum perturbation broadly across many distant residues, characteristic of residues that mediate long-range communication (allosteric hubs, hinge regions, or domain interface).

**FIG. 1:**
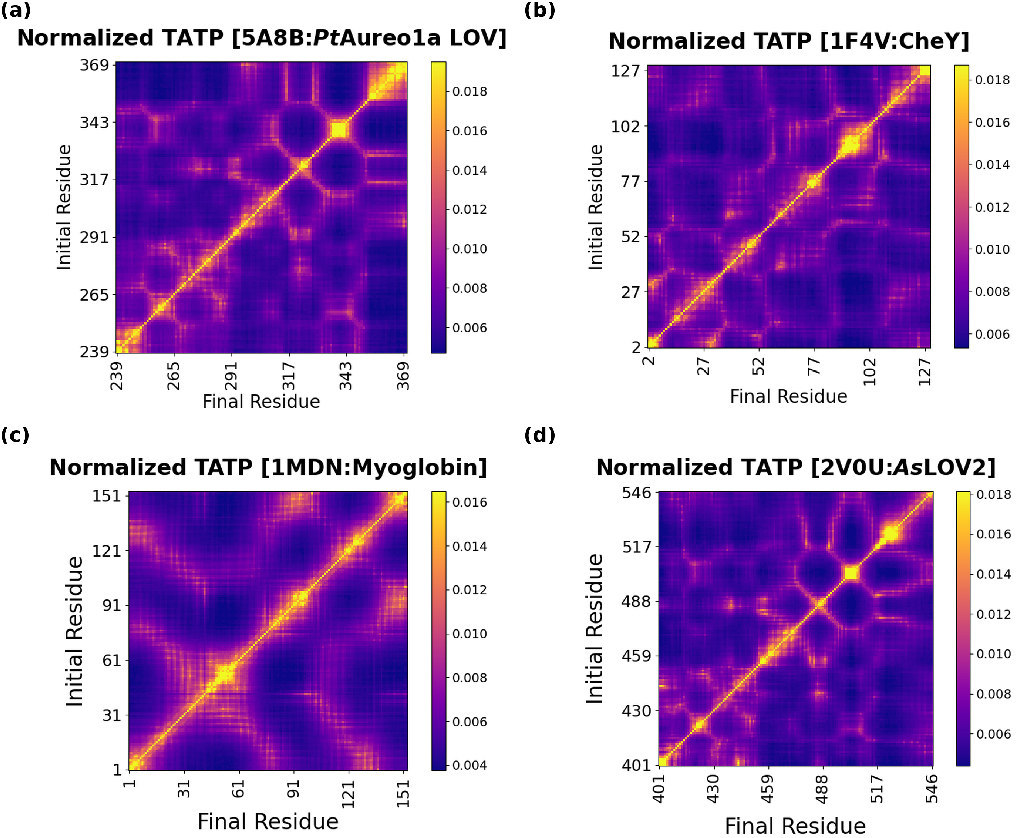
DTQW-derived normalized time-averaged transition probability (TATP) profiles for representative proteins: (a) *Pt*Aureo1a LOV, (b) CheY, (c) Myoglobin and (d) *As*LOV2.

Entropy is therefore an effective *perturbation-flow descriptor* that quantifies the degree of dynamical delocalization initiated from each residue. Higher entropy residues act as broad communication hubs whose activation or mutation would be expected to propagate effects across larger portions of the protein, explaining why such residues are often targets for disease-causing mutations and allosteric regulation.

FIG. 2 displays the per-residue normalized Shannon entropy (normalized values 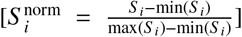 lie in the interval [0,1]) for the four representative proteins.

To identify statistically significant high-entropy residues, we computed a per-protein threshold based on the normalized entropy distribution:

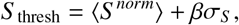

where ⟨*S* ^*norm*^⟩ is the mean entropy, *σ*_*S*_ is the standard deviation, and *β* is a multiplier (typically *β* = 0.7) chosen to identify residues in the upper tail 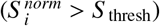 of the distribution.

c *Step 3: Identification of Contiguous High-Entropy Regions (CHERs)*. We identified contiguous stretches of residues with Shannon entropy above the per-protein statistical significance threshold as highlighted in FIG. 2 of the four systems with shaded regions. We have given more importance to the spatially clustered regions of high-entropy residues rather than scattered individual high-entropy residues for our framework. These contiguous high-entropy regions (CHERs) represent domains of residues which have more diversity and uncertainty of transitions from each residue of those domains to all other residues suggesting that these regions may have connection with the residues of long-range interaction. For our work, we would use top k most populated CHERs (with *k* = 3-4) for simplicity.
d *Step 4: The most probable inter-region transitions*. After identifying top k most populated CHERs for a protein, we checked the normalized transition probabilities between two distinct CHERs (inter-CHER transitions) and picked out top l most higher probabilistic transitions(with *l* = 3-4) and their corresponding residues. Then, all the residues collecting from all the different inter-CHER combination are identified as critical residues involved in long-range interactions of that protein.

**FIG. 2:**
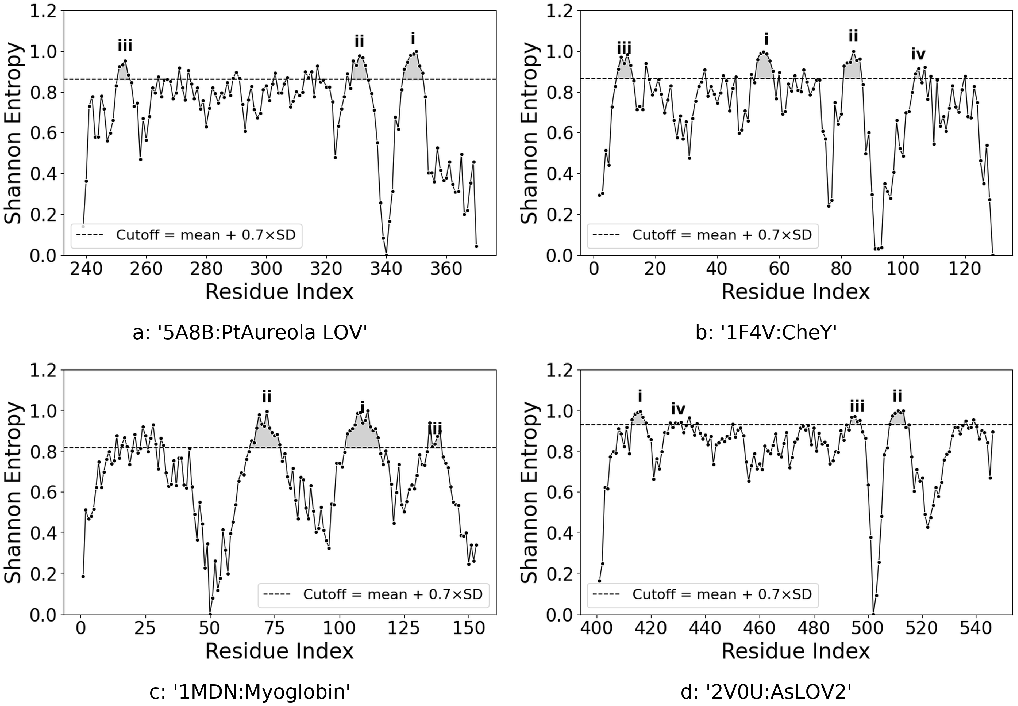
DTQW-derived entropy profiles for representative proteins: (a) *Pt*Aureo1a LOV, (b) CheY, (c) Myoglobin and (d) *As*LOV2. Shaded regions denote top k populated contiguous high-entropy regions (CHERs) exceeding the statistical cutoff.

**FIG. 3:**
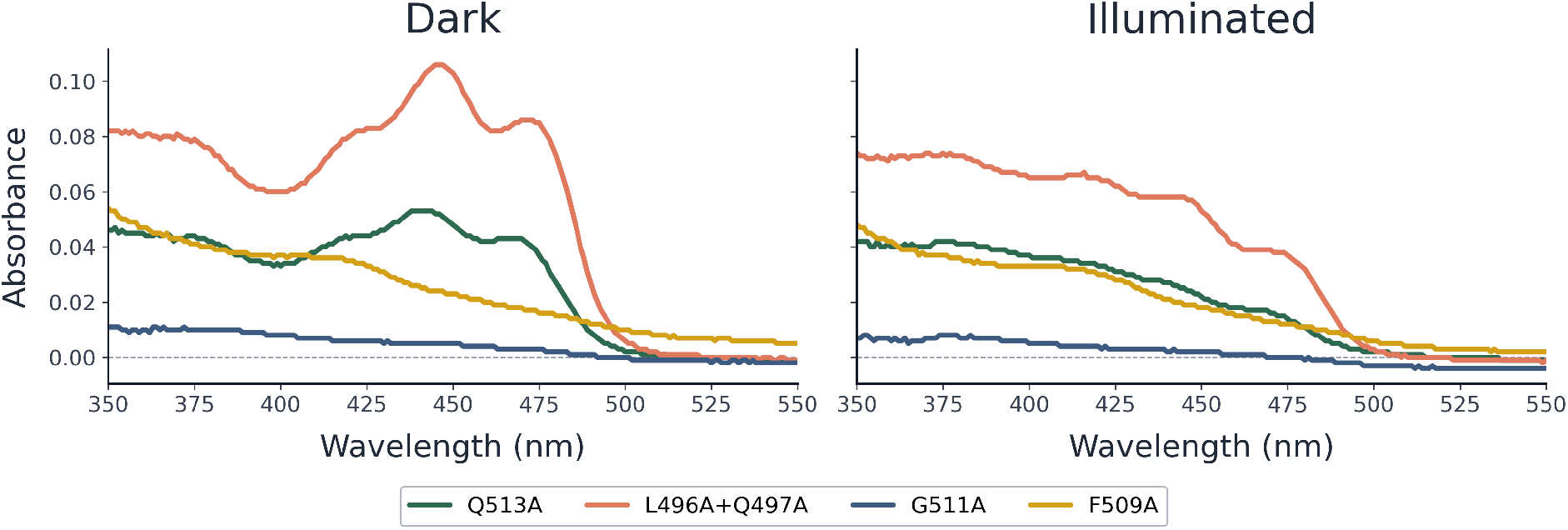
Dark and light state UV-Visible spectrum of wild-type *As*LOV2 and its mutants

Suppose, there are three top k most populated CHERs (with *l* = 3-4), denoted by *R*_*a*_, *R*_*b*_, and *R*_*c*_, in a protein. Then, a total of three distinct inter-CHER combinations are possible:

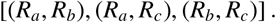

For each inter-CHER pair, we consider the corresponding transition set

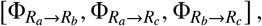

where each inter-CHER transition set is the set of all transitions between these two distinct pair, i.e.,

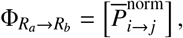

where *i* ∈ *R*_*a*_, *j* ∈ *R*_*b*_ are residue indices of those two distinct CHERs *R*_*a*_ and *R*_*b*_, respectively.

Then, we recorded the top l most probable transitions (with *l* = 3-4) from each set and got the critical residues according to our formalism.

Residue pairs exhibiting strong inter-CHER region transition probabilities are like *communication bridges* and mutations or conformational changes of those residues would most directly disrupt long-range interaction and communication signal propagation between distant functional sites. These bridges represent non-local couplings that are not obvious from three-dimensional proximity alone and highlight subtle structural pathways that evolution has exploited to enable functional regulation, shown in Table I .

#### 3. Correspondence of predicted residues with known functional sites

In all tested systems, the residues identified by our framework showed strong correspondence with experimentally and computationally established functional sites. A large fraction of the predicted residues coincide with known allosteric sites, or with residues lying on allosteric communication pathways previously identified by structural and biochemical studies [4, 8], normal mode analysis [34], crystallographic ensemble refinement [35], and molecular dynamics simulations [36]. This agreement indicates that broad, high-entropy quantum-walk spreading captures the same long-range couplings that these independent methods attribute to allosteric signalling.

The predicted residues are enriched at domain boundaries and inter-domain interfaces, and in segments flanking conformational hinges and flexible regions. Such regions are known to govern the relative motions of structural domains [37], and are therefore natural relay points for transmitting perturbations between distant parts of a protein. In the LOV photoreceptors AsLOV2 (PDB: 2V0U) [12] and aureochrome 1a from *Phaeodactylum tricornutum* (PDB: 5A8B) [38], for example, the identified residues lie on the *β*-scaffold strands that face the flanking helices, where light-induced structural changes are transmitted from the flavin chromophore to the terminal helices.

In enzymatic systems, the framework recovers catalytic residues together with their surrounding coordination spheres [39], and in regulatory proteins it identifies residues lining cofactor- and effector-binding pockets. In the prokaryotic (6-4) photolyase PhrB from *Agrobacterium tumefaciens* (PDB: 4DJA) [40], the (6-4) photolyase from *Caulobacter crescentus* (PDB: 9HNM) [41], cryptochrome B from *Rhodobacter sphaeroides* (PDB: 3ZXS) [42], and the animal-like cryptochrome from *Chlamydomonas reinhardtii* (CraCRY, PDB: 8Z1J) [43], the predicted residues map to the catalytic *α*-helical domain that harbours the FAD cofactor. In the response regulator CheY (PDB: 1F4V) [44], they include the phosphorylation site Asp57, the metal-coordinating Asp12, and the *β*4–*α*4 region involved in propagating the activation signal.

Finally, the identified residues frequently encompass ligand-binding interfaces, DNA-binding regions, and protein– protein interaction surfaces in regulatory proteins, as well as subunit interfaces in oligomeric proteins. These are well-known hotspots in which a small number of residues contribute disproportionately to binding and function [45, 46]. Examples include the dimerization surface of the aureochrome 1a LOV domain, which couples light sensing to DNA binding [38], and the FliM-binding surface of CheY [44].

### B. System-wise study: Applications and Validation

1F4V is the crystal structure of 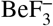 -activated *E. coli* CheY bound to the N-terminal peptide of the flagellar switch protein FliM [44]. Our approach identifies the acidic activesite residues ASP12 and ASP57, where phosphorylation by CheA initiates the signal [47, 48], together with TRP58. LEU81–VAL86 lie in the *β*4–*α*4 region next to the transmission residue THR87 [49]. SER104 and GLY105 flank TYR106, whose “in” rotamer creates the FliM binding surface [44, 50].

2V0U is the dark-state structure of the *Avena sativa* phototropin 1 LOV2 domain [12]. Blue light forms a CYS450– FMN adduct [17, 51], which is sensed by the conserved GLN513, a residue our approach captures [22]. PHE494, HIS495 and GLN497 relay the signal across the *β*-sheet, while ILE510 and VAL512 form the J*α* docking surface [12, 20]. ASN414 and ILE417/THR418 in the N-terminal flank mark the ASN414–ASP515 hydrogen bond that breaks on illumination [12].

1MDN is the structure of CO-bound pig myoglobin [52]. Ligand binding at the heme is controlled mainly by the distal HIS64 (E7) and VAL68 (E11), together with LEU29 (B10) and PHE43 (CD1), which shape the distal pocket [52, 53]. The proximal HIS93 (F8) links the heme iron to the protein, transmitting ligand-induced changes to the globin fold [53, 54].

5A8B is the dark-state, native-like parallel dimer of the *Phaeodactylum tricornutum* aureochrome 1a LOV domain [38, 55]. Blue light forms a cysteinyl–FMN adduct, and the signal is relayed through the LOV core to the flanking A^′^*α* and J*α* helices, which form the dimer interface [55–57]. Our approach identifies contiguous clusters at the N-terminal segment (ASN251–VAL254), the core (GLN291, ASN329, GLN330) and the C-terminal segment (GLN350–LYS352). These are consistent with light-induced unfolding of the flanking helices that drives dimerization and DNA binding [57, 58].

#### 1. Mutational Analysis of QW-Predicted Residues in AsLOV2

ILE417, THR418, LEU496, GLN497, TYR508, PHE509, ILE510 and GLY511 have been identified via theoretical predictions in *Avena sativa* Phot1 LOV2 (*As*LOV2) photosensor domain. Since this is a photosensor, the immediate connection that comes to mind is that the predicted residues must be related to light-dark photocycle kinetics. However, it is not necessary that mutating the identified residues would change the photocycle kinetics significantly. For example, the flipping GLN513 of *As*LOV2 has been the focal point of several seminal research articles [22, 59–61] But when it comes to dark recovery kinetics, mutation of GLN513 to ALA [62, 63] (in our experiment as positive control or Zayner et al., 2012) resulted in slightly slower dark recovery rate (8.4 × 10^−3^ sec^−1^ as measured in our study or a lifetime equal to 261 seconds according to Zayner et al. as compared to 81 seconds lifetime as measured for wild type). As determined by Zayner et al., most of the mutations barring ASN414 to VAL, resulted in almost unchanged dark recovery kinetics in *As*LOV2. Therefore, besides conducting dark recovery kinetics on the mutants of the mathematically predicted residues, we carefully observe the PDB structure and try to validate our findings. The detailed importance of the residues has been mentioned in Table II.

**TABLE II:**
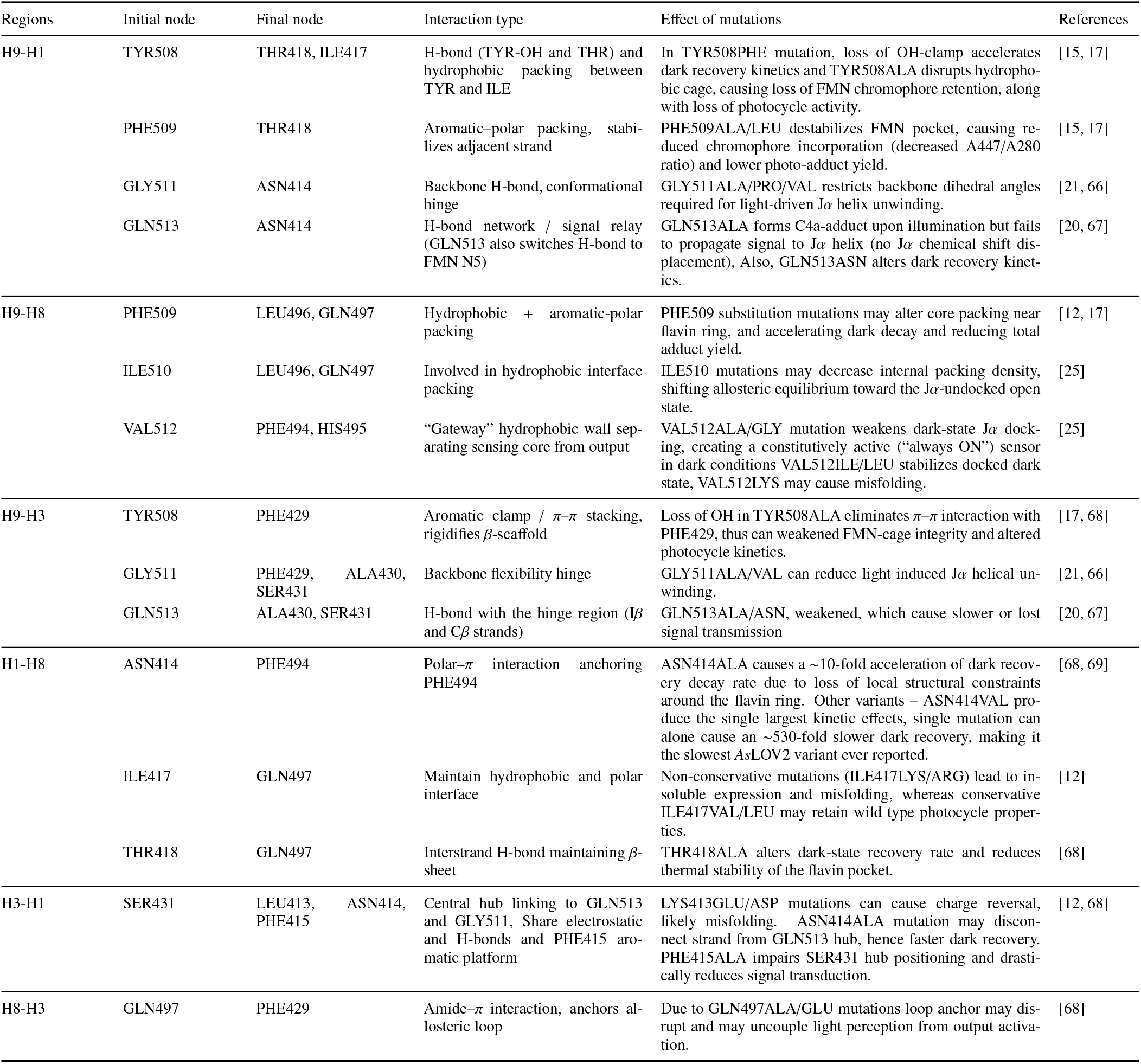
Predicted residue interactions and effects of mutations in *As*LOV2.

| Regions | Initial node | Final node | Interaction type | Effect of mutations | References |
| --- | --- | --- | --- | --- | --- |
| H9-H1 | TYR508 | THR418, ILE417 | H-bond (TYR-OH and THR) and hydrophobic packing between TYR and ILE | In TYR508PHE mutation, loss of OH-clamp accelerates dark recovery kinetics and TYR508ALA disrupts hydrophobic cage, causing loss of FMN chromophore retention, along with loss of photocycle activity. | [15, 17] |
|  | PHE509 | THR418 | Aromatic-polar packing, stabilizes adjacent strand | PHE509ALA/LEU destabilizes FMN pocket, causing reduced chromophore incorporation (decreased A447/A280 ratio) and lower photo-adduct yield. | [15, 17] |
| | GLY511 | ASN414 | Backbone H-bond, conformational hinge | GLY511ALA/PRO/VAL restricts backbone dihedral angles required for light-driven $J\alpha$ helix unwinding. | [21, 66] |
| | GLN513 | ASN414 | H-bond network / signal relay (GLN513 also switches H-bond to FMN N5) | GLN513ALA forms C4a-adduct upon illumination but fails to propagate signal to $J\alpha$ helix (no $J\alpha$ chemical shift displacement), Also, GLN513ASN alters dark recovery kinetics. | [20, 67] |
| H9-H8 | PHE509 | LEU496, GLN497 | Hydrophobic + aromatic-polar packing | PHE509 substitution mutations may alter core packing near flavin ring, and accelerating dark decay and reducing total adduct yield. | [12, 17] |
| | ILE510 | LEU496, GLN497 | Involved in hydrophobic interface packing | ILE510 mutations may decrease internal packing density, shifting allosteric equilibrium toward the $J\alpha$-undocked open state. | [25] |
| | VAL512 | PHE494, HIS495 | “Gateway” hydrophobic wall separating sensing core from output | VAL512ALA/GLY mutation weakens dark-state $J\alpha$ docking, creating a constitutively active (“always ON”) sensor in dark conditions VAL512ILE/LEU stabilizes docked dark state, VAL512LYS may cause misfolding. | [25] |
| H9-H3 | TYR508 | PHE429 | Aromatic clamp / π-π stacking, rigidifies β-scaffold | Loss of OH in TYR508ALA eliminates π-π interaction with PHE429, thus can weakened FMN-cage integrity and altered photocycle kinetics. | [17, 68] |
| | GLY511 | PHE429, ALA430, SER431 | Backbone flexibility hinge | GLY511ALA/VAL can reduce light induced $J\alpha$ helical unwinding. | [21, 66] |
| | GLN513 | ALA430, SER431 | H-bond with the hinge region ($I\beta$ and $C\beta$ strands) | GLN513ALA/ASN, weakened, which cause slower or lost signal transmission | [20, 67] |
| H1-H8 | ASN414 | PHE494 | Polar-π interaction anchoring PHE494 | ASN414ALA causes a $\sim 10$-fold acceleration of dark recovery decay rate due to loss of local structural constraints around the flavin ring. Other variants – ASN414VAL produce the single largest kinetic effects, single mutation can alone cause an $\sim 530$-fold slower dark recovery, making it the slowest AsLOV2 variant ever reported. | [68, 69] |
|  | ILE417 | GLN497 | Maintain hydrophobic and polar interface | Non-conservative mutations (ILE417LYS/ARG) lead to insoluble expression and misfolding, whereas conservative ILE417VAL/LEU may retain wild type photocycle properties. | [12] |
| | THR418 | GLN497 | Interstrand H-bond maintaining β-sheet | THR418ALA alters dark-state recovery rate and reduces thermal stability of the flavin pocket. | [68] |
| H3-H1 | SER431 | LEU413, ASN414, PHE415 | Central hub linking to GLN513 and GLY511, Share electrostatic and H-bonds and PHE415 aromatic platform | LYS413GLU/ASP mutations can cause charge reversal, likely misfolding. ASN414ALA mutation may disconnect strand from GLN513 hub, hence faster dark recovery. PHE415ALA impairs SER431 hub positioning and drastically reduces signal transduction. | [12, 68] |
| H8-H3 | GLN497 | PHE429 | Amide-π interaction, anchors allosteric loop | Due to GLN497ALA/GLU mutations loop anchor may disrupt and may uncouple light perception from output activation. | [68] |

In the case of ILE417 and THR418, ILE417 is located at the critical juncture of TYR508, LEU546 and LEU408 in close proximity thereby making hydrophobic contacts. It is a well-established fact that *As*LOV2 is the most commonly adopted scaffold to be used in LOV-based optogenetics designs, and the molecular basis for this is the efficient folding and unfolding of the terminal J*α* and A^′^-*α* helices onto and from the *β*-core, respectively, in the dark and light states. ILE417 is indeed strategically placed to make hydrophobic contacts with TYR508 from A^′^-*α* and LEU546 from J*α* helices, respectively—essential to maintain “off state” under dark conditions. Mutation to longer or shorter chain amino acids could be inappropriate to maintain hydrophobic contacts with precision lengths. A recent study on mutation of the homologous ILE414 in ToAureo1, selected through random mutagenesis [64], abrogated flavin-binding activity. THR418 is making a two-pronged contact—while the nonpolar tip is contacting FMN’s C7, the polar tip is making contacts with residues from I*β* (PHE509, also identified in this study) and the loop connecting A*β* and B*β*. A mutation in homologous CYS76 to ALA/VAL prolongs dark recovery rate in VVD photoreceptor [65].

LEU496 and GLN497 are both located at G*β* (G-*β*) in the *As*LOV2 core. Mutation of both LEU496 and GLN497 to Ala does not significantly alter the photocycle kinetic behavior (2.2 × 10^−2^ sec^−1^ as determined by this study compared to 1.3 × 10^−2^ sec^−1^ for WT as determined by [65]). LEU496 is among the few “inward facing” hydrophobic residues surrounding the FMN binding pocket. In NcVVD, the homologous residue of LEU496 is MET165, mutation of which leads to altered steric and electronic environment at the flavin binding site and thus influences dark recovery kinetics. While LEU496 makes important hydrophobic contact with flavin and PHE509, GLN497 makes polar contact with adjacent HIS495 and neighbouring LYS533 from J*α* helix. Further, TYR508 is also engaged in making polar contact with the terminal residue of J*α* helix. Similarly, ILE510 also makes hydrophobic contact with J*α* residues, ILE539 and ASP540. GLY511 mutation to MET led to failure to procure holoprotein [65], showing the requirement of a smaller amino acid in this particular location.

Overall, our study identifies amino acid residues, often adjacent in primary structure, from the single state structure of *As*LOV2 [PDB ID: 2V0U]. In most cases, while one residue makes an important contact with the active site flavin, the adjacent one ensures binding with the critical A^′^*α* or J*α* helix. Simultaneous preservation of the steric and electronic environment of the flavin while maintaining crucial contacts with the terminal helices makes *As*LOV2 a functionally efficient photosensor. Our study essentially captures these critical junctures.

## III. CONCLUSION

We have introduced a quantum walk and entropy-based computational framework for identifying functionally important residues mediating long-range interactions and communication in proteins. By applying discrete-time quantum walk (DTQW) dynamics to protein contact networks (PCNs) derived from crystallographic structures, we capture coherent, interference-driven information propagation and get biologically significant residues analysing the transition probabilities using Shannon entropy.

The framework has three basic grounds:

1. **Quantum coherence and interference:** DTQW dynamics maintains coherence and naturally incorporates superposition and interference. It enables the walker to explore multiple paths simultaneously on the network revealing structured transport patterns unlike classical diffusion.
2. **Shannon entropy quantification:** High entropy of the normalized TATP distribution identifies residues from which perturbations spread broadly across the protein, serving as effective communication hubs of long-range interaction and communication.
3. **Structure-only framework:** The approach requires only three-dimensional structural coordinates and a standard contact definition (4.5 Å). So, it does not depend on molecular dynamics simulations, sequence alignments, or prior functional data.

To demonstrate broad applicability and generalizability, we chose diverse category of 8 proteins. We have identified residues involved in high-probability inter-CHER transitions show strong correspondence with experimentally validated functionally important residues involved in long-range interactions and communications expressed differently as allosteric sites, catalytic residues, and regulatory hotspots etc. Our framework thus establishes a new paradigm for computational structural biology, leveraging quantum mechanical principles to predict functionally critical residues with a good accuracy and interpretability. This framework might have been used in many areas i.e., drug discovery, protein engineering, disease research, systems biology etc. for preliminary guess of critical residues involved in long-range interactions and understanding before going into the actual experiments. Future extensions could incorporate ensemble-based approaches using multiple conformations from molecular dynamics or cryo-EM ensembles, weighted DTQW formulations incorporating different types of initial states. The theoretical foundation of this framework rooted in/grounded on established quantum walk theory and information theory. So, it provides a robust basis for continued development and application across diverse protein systems and biological networks.

## IV. MATERIALS AND METHODS

### A. Protein Contact Network (PCN) Construction

Protein three-dimensional structures were obtained from the Protein Data Bank (PDB) and converted into residue-level protein contact networks (PCN). Each amino acid residue is represented as a node, and an undirected edge is formed between two residues if any pair of their atoms lie within a cutoff distance of 4.5 Å. This cutoff captures non-bonded interactions (van der Waals contacts, hydrogen bonds, and electrostatic interactions) that stabilize the tertiary structure and responsible for many basic functions.

The resulting network is represented by an undirected adjacency matrix **A** of dimension *N* × *N*, where *N* is the number of residues:

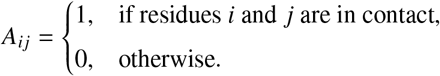

Self-loops (diagonal elements) are excluded. The degree of residue *i* is *d*_*i*_ = ∑_*j*_ *A*_*i j*_. The degree matrix **D** is defined as a diagonal matrix with *D*_*ii*_ = *d*_*i*_. The graph Laplacian **L** = **D** − **A** is a fundamental matrix to define the whole characteristics of the network.

### B. Discrete-Time Quantum Walk (DTQW) Formalism

#### 1. Hilbert Space and Initial State

In a discrete-time quantum walk, the Hilbert space is a tensor product of the position space *H*_*P*_ spanned by the network nodes {|*i* ⟩: *i* = 1, …, *N*} and a coin space *H*_*C*_ encoding maximum possible local outgoing directions at each individual node:

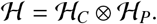

The walker is initialized at residue *i*_0_ in a uniform mixture over all its outgoing coin states. The corresponding initial density matrix is [70]

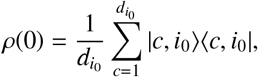

where 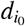 is the degree (number of outgoing edges) of node *i*_0_ and c labels the coin (direction) index.

#### 2. Unitary Evolution and Coin Operators

Each step of evolution is guided/defined by a unitary coin operator **C** followed by a shift operator **S** that moves probability amplitudes between connected residues:

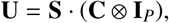

where the coin operator acts on the coin space only and **I**_*P*_ is the identity operator, acts on the position space only. Each outgoing edge |*c, i*_0_ ⟩ evolves independently under the unitary operator **U**, and the overall evolution of the mixed state is obtained by averaging over all such evolutions according to the initial state.

The coin operators are of many different types of different characters i.e., Fourier coin, Grover coin etc., where

##### Fourier coin

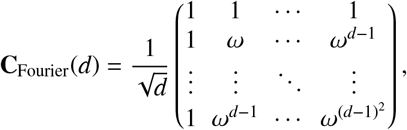

where *ω* = *e*^2*πi/d*^ is a primitive *d*-th root of unity. This coin produces symmetric, broad propagation.

##### Grover coin

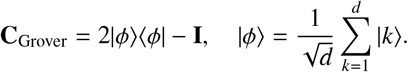

This coin produces narrower, more localized spreading emphasizing nearby residues.

The shift operator **S** is graph-dependent and moves probability amplitudes along network edges:

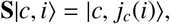

where *j*_*c*_(*i*) is the *c*-th neighbor of residue *i* in the PCN. This operation is unitary because the network graph is connected and undirected.

#### 3. Transition Probability Computation

At time *t*, the state is given by

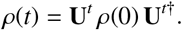

The instantaneous transition probability from initial residue *i*_0_ to target residue *j* at time *t* is then computed as

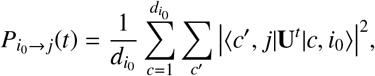

which corresponds to averaging the probabilities obtained by evolving each outgoing edge of *i*_0_ separately.

#### 4. Time-Averaged Transition Probability (TATP)

Due to quantum interference, 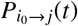 exhibits oscillations with time.To obtain a stationary communication measure, we compute the time-averaged transition probability (TATP) [70]:

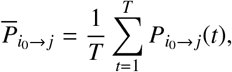

which captures long-time residue-to-residue communication likelihood by averaging over interference-induced fluctuations. We have used *T* = 100 steps. This choice of *T* was selected to balance convergence (steady-state behavior) with computational efficiency. For PCN networks (on an average 500 residues), 100 steps was empirically found to achieve stable TATP values.

The full TATP matrix is constructed by iterating the walk initiation over all *N* residues as starting points, producing an *N* × *N* matrix 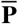 where entry (*i, j*) represents the time-average transition probability of reaching residue *j* starting from residue *i*. Because each time step of the DTQW is governed by a unitary operator, the total probability is conserved during evolution. After long-time averaging, the transition matrix 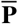 becomes approximately *doubly stochastic*, with both row and column sums close to one. This reflects a balanced bidirectional flow of probability across the network, signifying that, on average, each residue both sends and receives comparable quantum information over time.

### C. Shannon Entropy Calculation

The Shannon entropy related to probability distribution is computed as

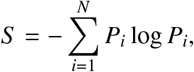

using natural logarithm (ln).

#### Construction of *As*LOV2 mutants by site-directed mutagenesis

The plasmid pET-28a(+) encoding wild-type oat *As*LOV2 residues (UniProt: O49003) is used for the experiments. This is an N-terminal hexahistidine (His6), which is used as the template for all site-directed mutagenesis (SDMs) reactions. The identity of the wild-type construct was confirmed by bidirectional Sanger sequencing of the full coding insert prior to mutagenesis PCR. Codon-optimised wild-type sequence of *As*LOV2 with N-terminal hexaHistidine tag, *His*_6_ was used as a template for further SDMs.

Mutagenic oligonucleotide primers were designed for each of the four mutant constructs described in Table III. Each forward primer was designed to incorporate the desired codon change, with the mismatched nucleotide(s) positioned at the center of the primer. SDM PCR reactions contained 1 × Phusion GC master mix, 150–200 ng template, 0.5 *µ*M each primer and 3% (v/v) DMSO. The methylated parental template was removed by DpnI digestion (5 U, New England Biolabs), and the product was transformed into chemically competent *E. coli* DH5*α* with selection on LB agar containing 50 *µ*g/mL kanamycin. Plasmid DNA from individual colonies was verified by Sanger sequencing using T7 promoter and T7 terminator primers.

**TABLE III:**
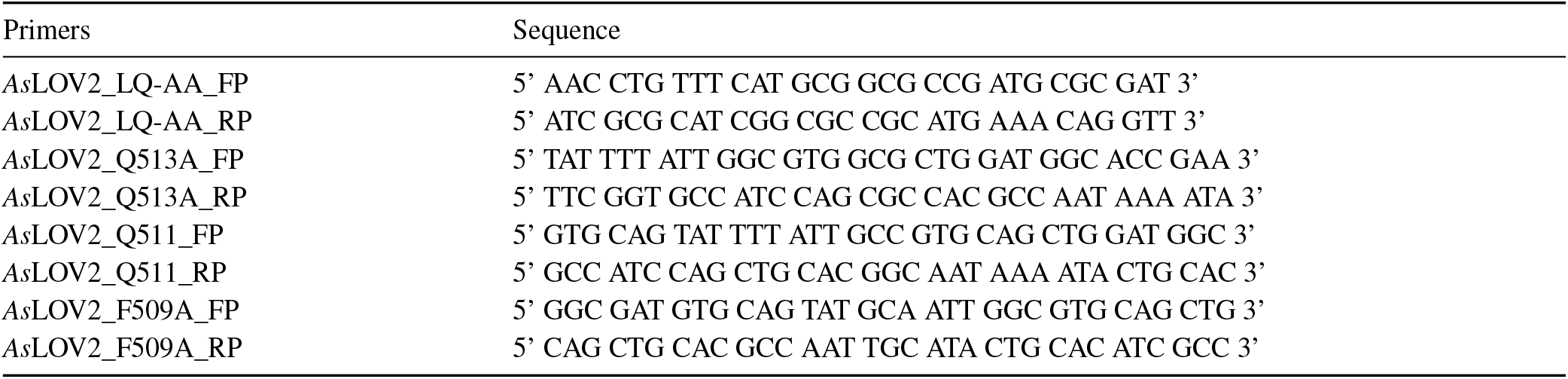
Primers.

#### Transformation and large-scale protein expression of mutants *As*LOV2 in *E. coli* C43(DE3)

Chemically competent *Escherichia coli* strain C43(DE3) was selected as the expression host for all *As*LOV2 constructs (mutants + wild type). These cells were transformed with 1 ng of each construct by the heat shock method at 42^°^C for 1 min 10 sec and recovered in 700 *µ*L Luria broth medium at 37^°^C for 1 hour and plated (Luria Agar) in the presence of Kanamycin (50 *µ*g/mL). Single colonies were used to inoculate 10 mL starter culture containing kanamycin and were grown overnight at 37^°^C at 150 rpm. The following day, 2 liters of LB broth was inoculated by 1% starter culture and incubated at 37^°^C at 150 rpm till O.D. reached ∼ 0.6. The protein was induced in the presence of 300 *µ*M isopropyl *β*-D-thiogalactopyranoside (IPTG). Induced culture was kept at 22^°^C for 16 hours. Bacterial cells were then harvested by centrifugation at 8000 rpm for 10 minutes. Further, cell pellets were resuspended in lysis buffer (20 mM Tris, pH 8.0, 50 mM NaCl, 10% glycerol) followed by addition of lysozyme and Protease cocktail inhibitor. The treated cells were sonicated and centrifuged at 13,000 rpm for 1 hour at 4^°^C. The supernatant was incubated with FMN and pre-equilibrated Ni-NTA resins. The supernatant was passed through the empty Ni-NTA columns under gravity separation, and flow through was collected. The column was rinsed with wash buffer (20 mM Tris, pH 8.0, 50 mM NaCl, 10% glycerol, 10 mM imidazole). Freshly prepared wash buffer is then used to elute the desired protein fraction. Freshly prepared elution buffer is used to elute the desired protein fraction. Further, desalting/gel-filtration was performed to remove excess imidazole by using PD10 desalting columns after using the same equilibration buffer. For gel filtration chromatography, the protein was loaded onto a gel filtration column, and the desired protein fractions were pooled and used for dark recovery kinetics by using UV-Vis spectroscopy.

#### Dark recovery kinetics

Recovery kinetics were captured with UV-Vis spectroscopy of all the constructs after measuring concentrations of protein at O.D. at 447 nm in the dark state. Further, absorption spectra for all the constructs were recorded using a UV-Vis double-beam spectrophotometer at 280 nm in the dark state and after 10 mins of illumination with 364 W/m2 LED light. The O.D. at 447 nm values were plotted against time using an exponential equation [O.D. at 447nm = constant (1 − *e*^−*kt*^)] to calculate the rate constant (k) and lifetime of the constructs.

### D. Computational Efficiency and Platform

For typical system sizes(proteins of approx. upto *N* = 200 residues), the DTQW simulations were performed in Python 3.9 or 3.10 using Networkx, NumPy, SciPy. For a typical protein with *N* = 200 residues and average degree 3 (average edge count per residue), DTQW evolution for 100 time steps and all 200 initial residues required <120 CPU-minutes on a standard workstation (Intel i9, 16 GB RAM). For bigger systems, we have used paramrudra supercomputer at SNBNCBS and use mpi parallelism over multi nodes and multi cores as required. Parallelization over initial residues and time steps could reduce time and uses of multi cores and nodes give more access to RAM memory for calculations.

## E. Data Availability and Reproducibility

All PDB structures used are taken from publicly available the Protein Data Bank (www.rcsb.org).

## F. Acknowledgements

DM and SR gratefully acknowledge the financial support provided by the Science and Engineering Research Board (SERB), Department of Science and Technology (DST), Government of India, under Sanction Order No. CRG/2023/006463. We thank Faruque Siddique for helpful discussions.

## Notes

### Competing Interest Statement

The authors have declared no competing interest.

